# The influence of the way of regression on the results obtained by the receptorial responsiveness method (RRM), a procedure to estimate a change in the concentration of a pharmacological agonist near the receptor

**DOI:** 10.1101/2023.06.23.546208

**Authors:** Ignac Ovari, Gabor Viczjan, Tamas Erdei, Barbara Takacs, Reka Szekeres, Nora Lampe, Judit Zsuga, Miklos Szucs, Zoltan Szilvassy, Bela Juhasz, Rudolf Gesztelyi

## Abstract

The receptorial responsiveness method (RRM) enables the estimation of a change in the concentration of a degradable agonist, near its receptor, by fitting its model to (at least) two concentration-effect (E/c) curves of a stable agonist of this receptor. One curve should be generated before this change in concentration, while the other one after this change, in the same (or in identical) biological system(s). It follows that RRM yields a surrogate parameter (“c_x_”), the concentration of the stable agonist that is equieffective with the change in the concentration of the degradable agonist. However, the curve fitting can be implemented several ways, which can affect accuracy, precision and convenience of use.

This study utilized data of previous *ex vivo* investigations. Known concentrations of stable agonists were estimated with RRM by performing individual (local) or global fitting (with one or two model(s)), combined with the use of a logarithmic (logc_x_) or non-logarithmic parameter (c_x_), and with ordinary least-squares or robust regression.

We found that the individual regression, the most complicated option, was the most accurate, followed closely by the moderately complicated two-model global regression and then by the easy-to-perform one-model global regression. The two-model global fitting was the most precise, followed by the individual fitting (closely) and by the one-model global fitting (from afar). The use of c_x_ and robust regression did not, whereas pairwise fitting (i.e. fitting only two E/c curves at once) did improve the quality of estimation. Thus, the two-model global fitting, performed pairwise, is recommended for RRM, but the individual fitting is a good alternative.

## 1. Introduction

Measuring the concentration of small molecules in biological samples is generally considered to be an issue for analytical chemistry that can now be performed to a high standard (Valcárcel, 2000). However, concentration determination of small molecules with a short half-life and/or strong compartmentalization tendency in living, functioning, moreover moving tissues is still a challenge (Karsai et al, 2006; Ramakers et al, 2008). If certain requirements are met, the receptorial responsiveness method (RRM) can help to solve this problem. RRM can quantify an acute (and occasionally chronic) increase in the concentration of a pharmacological agonist in the microenvironment of its receptors (Gesztelyi et al, 2004; Karsai et al, 2006).

To understand the principle of RRM (which may seem to be complicated at first glance, but is in fact simple), it is worth first considering a thought experiment. Let us administer the same dose of an agonist twice to a biological system, and then let us compute the effect of both doses. The correct way is to define the same initial state for both effects (the state before the administration of the first dose) and to take also the first dose into account when calculating the effect of the second one. However, if we define the state developed after the first dose as the initial state and ignore the first dose when computing the effect of the second dose, then we will get a smaller effect for the second dose than for the first one, although the two doses equal.

The explanation of this smaller effect is that the agonist molecules administered with the first dose consume part of the response capacity of the system by binding to a fraction of the receptors and then by eliciting an effect on them. For this reason, the agonist molecules administered with the second dose will act in a system with submaximal response capacity. If, for example, the dose in question is high enough to exert a (practically) maximal effect after its first administration, no further effect can develop, thus, when administering it for the second time, a system unable to respond will be experienced. Thus, this (experienced) decrease in the responsiveness depends on the magnitude of the ignored dose, more precisely on the agonist concentration near the relevant receptors. This phenomenon can be exploited to quantify an unknown agonist concentration developed in the tissue compartment of the binding site of the specific receptors (Gesztelyi et al, 2004; Grenczer et al, 2010a; Karsai et al, 2006, 2007).

It should be emphasized that this phenomenon, although superficially reminiscent of it, is not based on receptor desensitization, when the effect is correctly computed but smaller than expected. In turn, the receptor desensitization interferes with the above-mentioned phenomenon, thus this latter can only be used for concentration estimation if the receptor in question desensitizes slowly relative to the time window of the determination (about 10-40 min, depending on the circumstances). The A_1_ adenosine receptor belongs to the especially slowly desensitizing receptors (Willems et al, 2006; Klaasse et al, 2008; Mundell and Kelly, 2011).

The phenomenon described above can also be observed if the agonist being present first is different from that is added subsequently, but they evoke the same effect (so the same response capacity is consumed by them). For this, it is enough if they are agonists on the same receptor, or even if they influence the same postreceptorial signaling in the same direction (although binding to different receptors). The agonist that enters the biological system first and is later disregarded can be called as “biasing agonist”, and its concentration can be referred to as “biasing concentration” (Gesztelyi et al, 2004; Grenczer et al, 2010a).

A complete concentration-effect (E/c) curve (ranging from ineffective to saturative concentrations) encompasses the information needed for this concentration estimation with high redundancy, and thereby offers the possibility of a more reliable estimation than a single concentration - effect pair does. Consequently, the best way to assess a change (“bias”) in a biological response is to analyze E/c curves. Two types of E/c curves are required: one holding information on the real (intact) relationship between agonist concentration and effect in the given system (serving as a benchmark), and another one informing about the extent of the bias and thereby about the magnitude of the biasing concentration. These two E/c curves can be referred to as “intact” and “biased” that contain “intact” and “biased” effects, respectively. The corresponding intact and biased effects are evoked by the same agonist concentration under the same conditions, the only difference between them is the presence and neglect of a biasing concentration in the case of the biased effect (Gesztelyi et al, 2004; Grenczer et al, 2010a). It is worth anticipating that if more than one biasing concentration are to be estimated in the same system, then more than two E/c curves are to be assessed with RRM. This situation can be handled in two different (although theoretically similar) ways (see further on).

The essence of RRM is to fit its model to a biased E/c curve, utilizing the information that is implied in the corresponding intact E/c curve. The model of RRM describes the relationship between an intact effect and its biased counterpart by means of a quantitative receptor function model (e.g. the Hill equation: Gesztelyi et al, 2012). The information in the intact E/c curve can be made accessible two main ways. On one hand, the given receptor function model can be fitted to the intact E/c curve, and then the identical parameters in the RRM’s model can be constrained at the obtained values. After this “individualization”, the RRM’s model can simply be fitted to the corresponding biased E/c curve (Gesztelyi et al, 2004; Grenczer et al, 2010a, 2010b). In this paper (similarly to the recent ones: Szabo et al, 2019a; Viczjan et al, 2021, 2022), this procedure will be referred to as individual regression. This type of fitting may be classified as local regression, as the opposite of global regression (Anjos et al, 2020; Spatial Data Science, 2023). On the other hand, the intact and biased E/c curves can be fitted at once, *via* global regression. Global regression can also be performed in two ways: fitting the intact and biased E/c curves to one model (which is the model of RRM: Szabo et al, 2019a), or fitting these E/c curves to two models (the receptor function model and the compatible RRM’s model). This latter maneuver should be done so that the receptor function model fits only the intact E/c curve and the RRM’s model fits exclusively the biased E/c curve, with the common parameters shared between the two models (Curve Fitting Guide, 2023). Hereinafter, these two procedures will be referred to as one-model global regression and two-model global regression, respectively.

Anyway, RRM provides a measure of the biasing concentration as a best-fit value that can be a concentration (c_x_) or its logarithm (logc_x_). The nature of this measure depends on whether the biasing agonist and the agonist used for the E/c curves are the same or not. If yes, RRM directly estimates the biasing concentration (or its logarithm). If not, the best-fit value provided by RRM is the concentration (or its logarithm) of the agonist used for the E/c curves that is equieffective with the biasing concentration (Gesztelyi et al, 2004; Grenczer et al, 2010a). Although this latter estimate is just a surrogate parameter, it has the advantage that a stable agonist may be used to determine the biasing concentration of an agonist that is degradable and/or exhibits strong compartmentalization. As the estimate of RRM is derived from E/c curves, it applies to the (average) concentration of the biasing agonist near the receptors involved in the production of the effect measured (Gesztelyi et al, 2004; Karsai et al, 2006, 2007; Viczjan et al, 2022).

In a previous study (Szabo et al, 2019a), estimates of RRM were found to be more accurate and precise when obtained from individual regression rather than one-model global fitting. This is surprising in light of that the global regression is thought to be more powerful and reliable than the individual one, especially in the case of limited data and/or data with much scatter (Knutson et al, 1983; Motulsky and Christopoulos, 2004; Herman and Lee, 2012). The worse than expected performance of the one-model global fitting was attributed to the lack of an accurate estimate (c_x_) for the intact E/c curves, because the model of RRM, used in that study (Szabo et al, 2019a), contained logc_x_ as a parameter (instead of c_x_). Hence, RRM could not provide zero, the correct estimate for intact E/c curves, as the logarithm of zero is not defined.

Earlier, computer programs calculated only symmetrical confidence intervals (CIs). Therefore, quantities following log-normal distribution (e.g. concentrations) were recommended to be used as common logarithms in the equations to fit, in order to get correct CIs (Motulsky and Christopoulos, 2004). If asymmetry of CIs can be handled (not uncommon for modern software systems), the use of concentrations as a logarithm in the regression models seems to be less important. Thus, in the present study, one of our goals was to explore whether the model of RRM with c_x_ could be superior to that with logc_x_. Consistent with this, we aimed also to investigate whether the two-model global regression, another way to avoid fitting logc_x_ to an intact E/c curve, could provide better results that the one-model global regression, and perhaps than the individual fitting. In addition, issues of the distribution of the scatter of E/c curve data and of fitting E/c curve families (more than one biased E/c curve belonging to one intact E/c curve) were also addressed.

## 2. Materials and methods

### 2.1. Data analyzed

Investigations of the present study were carried out on cumulative, graded E/c curves, originally published in two earlier studies (Szabo et al, 2019a; Gesztelyi et al, 2004). The E/c curves were constructed with A_1_ adenosine receptor agonists, by measuring the contractile force of isolated, paced guinea pig left atria. These E/c curves, consisting of related intact and biased ones, were reevaluated with RRM, implemented by combining several ways of regression. For the intact - biased E/c curve pairs (obtained from Szabo et al, 2019a), using the RRM’s model containing logc_x_ *vs* c_x_, individual *vs* one-model global *vs* two-model global regression, and ordinary least-squares (simply: ordinary) *vs* robust fitting were combined. For the E/c curve families (obtained from Gesztelyi et al, 2004), individual *vs* one-model global *vs* two-model global fitting, ordinary *vs* robust fitting, furthermore fitting all related E/c curves at once *vs* fitting an intact E/c curve together with only one corresponding biased E/c curve at once were combined.

The intact E/c curves were generated with CPA, NECA or CHA, stable, synthetic A_1_ adenosine receptor agonists with long half-life (for more detail, see legends of Fig. 1-3), in the absence of any previously added adenosine receptor agonist. The biased E/c curves were constructed with the same agonists, but in the presence of a single known concentration of the agonist used for the E/c curve. This surplus agonist concentration was administered before the E/c curve (which latter was started when the effect of the former had fully developed) but it was disregarded during the evaluation (to make it a biasing concentration).

**Fig. 1.**
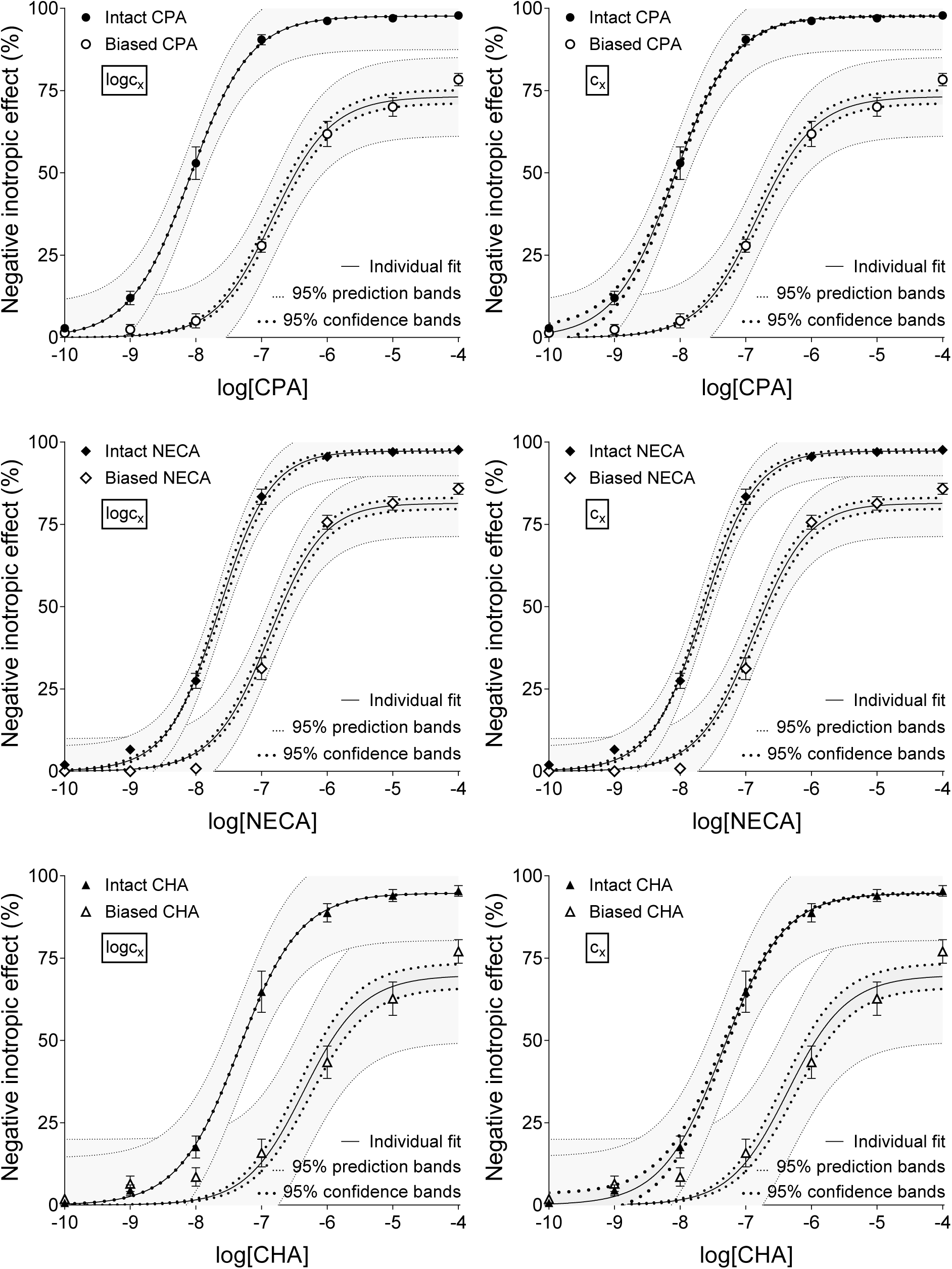
The implementation of RRM with individual regression combined with ordinary least-squares fitting, using two equations: one that contained the common logarithm of the concentration to be determined (logc_x_; left panels), and another one containing this concentration itself (c_x_; right panels), as a parameter. The measurement was performed on “Intact” – “Biased” concentration-effect (E/c) curve pairs obtained from a previous *ex vivo* study (Szabo et al, 2019a). The *x*-axis shows the common logarithm of the molar concentration of one of three synthetic A_1_ adenosine receptor full agonists (CPA, NECA, CHA), administered for the construction of the E/c curve. The *y*-axis shows the effect (the percentage decrease in the initial contractile force of isolated, paced guinea pig left atria). The “Intact” E/c curves (filled symbols) were generated in a conventional way, whereas the “Biased” E/c curves (open symbols) were constructed in the presence of a single extra concentration of the agonist used for the E/c curve. The symbols indicate the responses to the agonists averaged within the groups (± SEM). The continuous lines show the best-fit curves of the fitted equation 4 or 5 (containing logc_x_ or c_x_ as a parameter, respectively). The thick dotted lines indicate the 95% confidence bands, while the thin dotted lines denote the 95% prediction bands. RRM: receptorial responsiveness method; CPA: N^6^-cyclopentyladenosine; NECA: 5′-(N-ethylcarboxamido)adenosine; CHA: N^6^-cyclohexyladenosine

**Fig. 2.**
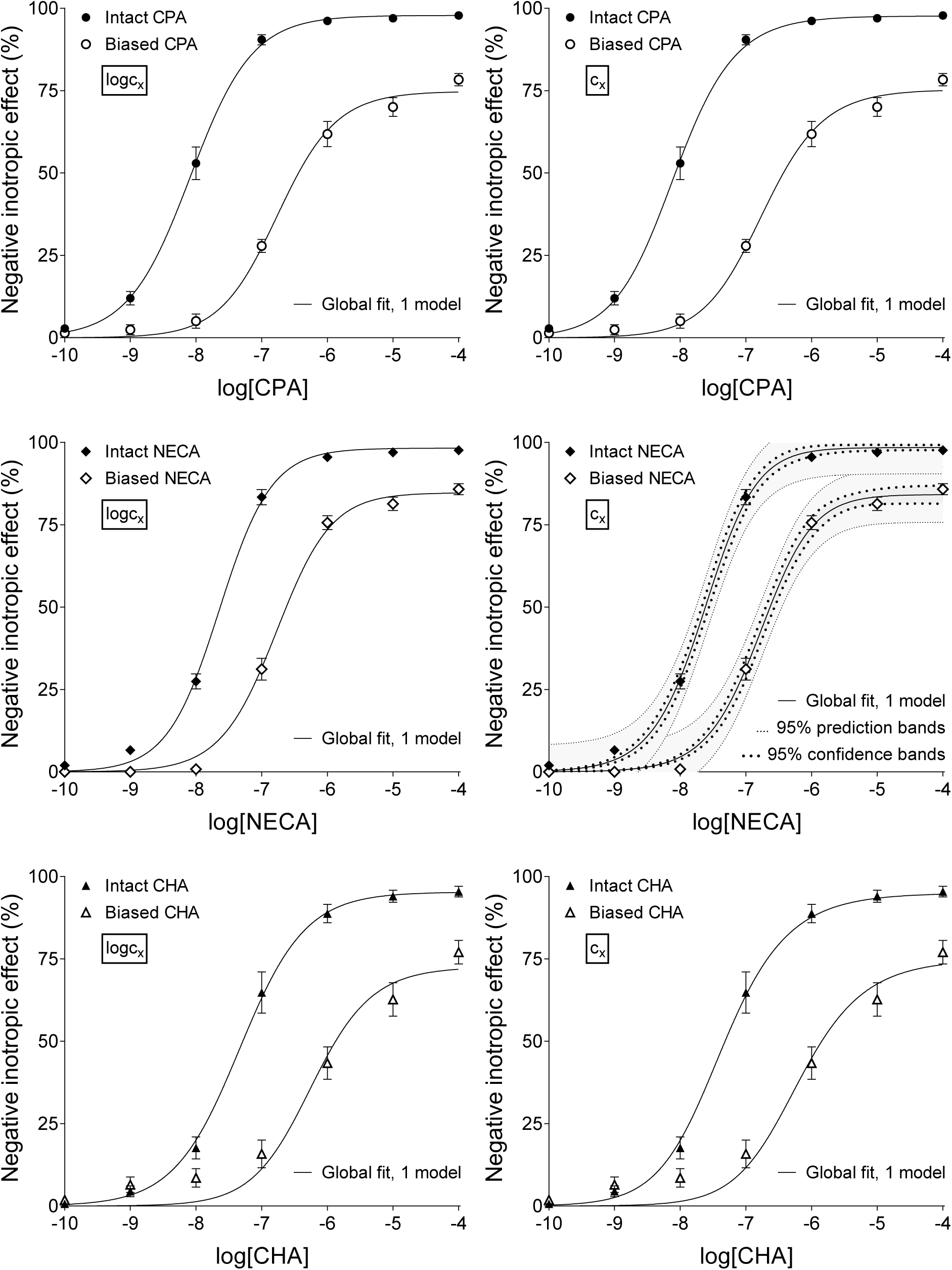
The implementation of RRM with one-model global regression combined with ordinary least-squares fitting, using two equations: one that contained the common logarithm of the concentration to be determined (logc_x_; left panels), and another one containing this concentration itself (c_x_; right panels), as a parameter. This procedure was performed on “Intact” – “Biased” concentration-effect (E/c) curve pairs obtained from an earlier *ex vivo* study (Szabo et al, 2019a). The *x*-axis shows the common logarithm of the molar concentration of one of three synthetic A_1_ adenosine receptor full agonists (CPA, NECA, CHA), administered for generating the E/c curve. The *y*-axis indicates the effect (the percentage decrease in the initial contractile force of isolated, paced guinea pig left atria). The “Intact” E/c curves (filled symbols) were constructed conventionally, whereas the “Biased” E/c curves (open symbols) were generated in the presence of a single surplus concentration of the agonist used for the E/c curve. The symbols indicate the responses to the agonists averaged within the groups (± SEM). The continuous lines show the best-fit curves of the fitted equation 4 or 5 (containing logc_x_ or c_x_ as a parameter, respectively). The thick dotted lines indicate the 95% confidence bands, while the thin dotted lines denote the 95% prediction bands (if any). RRM: receptorial responsiveness method; CPA: N^6^-cyclopentyladenosine; NECA: 5′-(N-ethylcarboxamido)adenosine; CHA: N^6^-cyclohexyladenosine

**Fig. 3.**
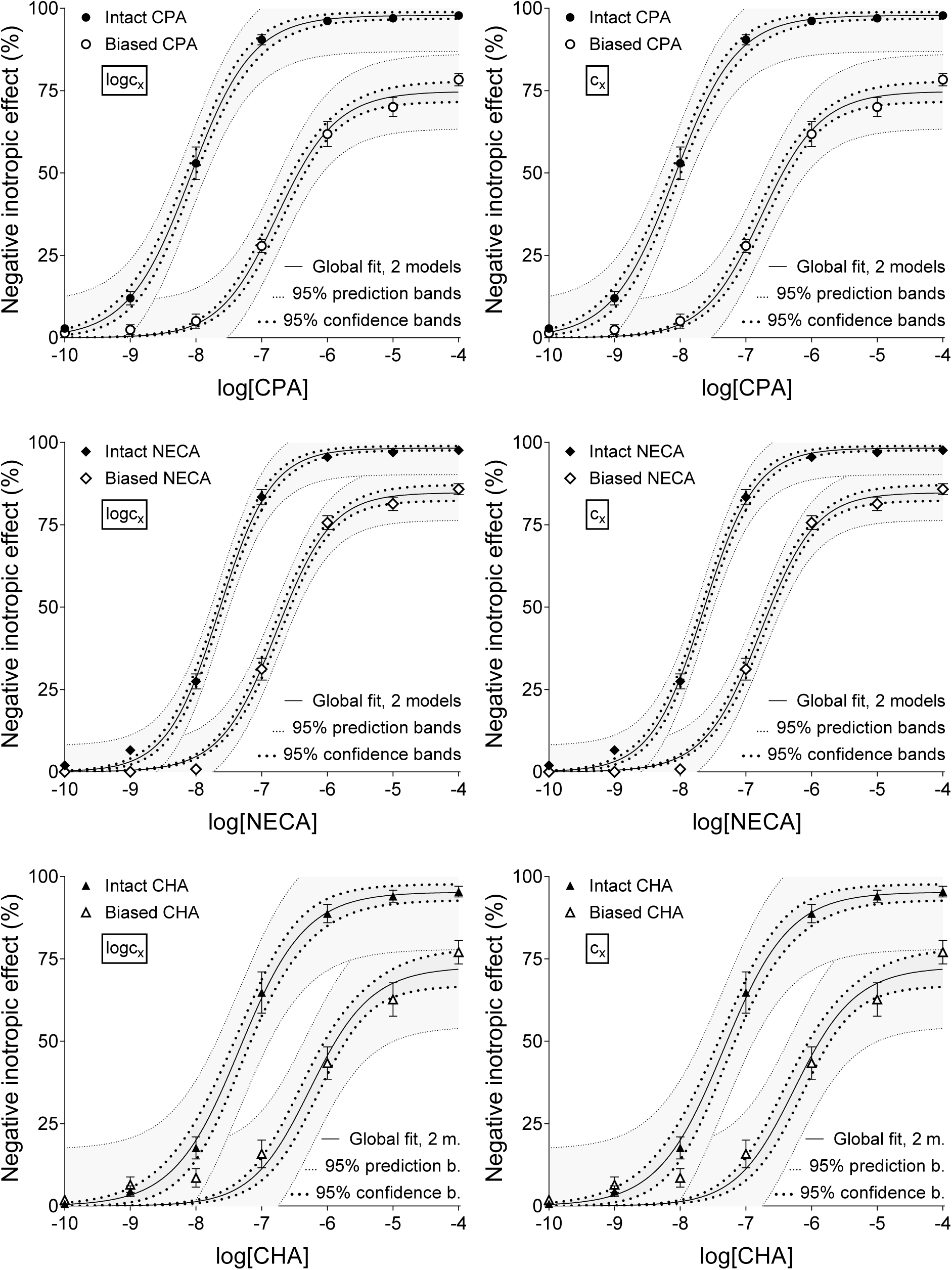
The implementation of RRM with two-model global regression combined with ordinary least-squares fitting, using two equations (in addition to the two models of the global fitting): one that contained the common logarithm of the concentration to be determined (logc_x_; left panels), and another one containing this concentration itself (c_x_; right panels), as a parameter. This procedure was performed on “Intact” – “Biased” concentration-effect (E/c) curve pairs obtained from a previous *ex vivo* study (Szabo et al, 2019a). The *x*-axis shows the common logarithm of the molar concentration of one of three synthetic A_1_ adenosine receptor full agonists (CPA, NECA, CHA), administered for constructing the E/c curve. The *y*-axis indicates the effect (the percentage decrease in the initial contractile force of isolated, paced guinea pig left atria). The “Intact” E/c curves (filled symbols) were generated in a conventional way, while the “Biased” E/c curves (open symbols) were constructed in the presence of a single extra concentration of the agonist used for the E/c curve. The symbols indicate the responses to the agonists averaged within the groups (± SEM). The continuous lines show the best-fit curves of the equation 2 fitted together with the equation 4 or 5 (containing logc_x_ or c_x_ as a parameter, respectively). The equation 2 was fitted to the “Intact” E/c curves, whereas the equation 4 (left panels) or 5 (right panels) was fitted to the “Biased” ones. The thick dotted lines indicate the 95% confidence bands, while the thin dotted lines represent the 95% prediction bands. RRM: receptorial responsiveness method; CPA: N^6^-cyclopentyladenosine; NECA: 5′-(N-ethylcarboxamido)adenosine; CHA: N^6^-cyclohexyladenosine

All E/c curves, used for the present study, were produced from individual E/c curves (each one belonging to one atrial preparation) through averaging them within the experimental groups by the curve fitting software in a way that individuality of the effect values (as “replicate Y values”) was preserved (Curve Fitting Guide, 2023). Each E/c curve mentioned in this work is the result of such a process.

One of our two experimental data sets (Szabo et al, 2019a) consisted of intact - biased E/c curve pairs (with biasing concentrations near to the EC_50_ for the agonist used), while the other one (Gesztelyi et al, 2004) contained three biased E/c curves (with biasing concentrations spanning two orders of magnitude including the EC_50_) to each intact E/c curve.

### 2.2. Regression manners dealing with models, variable parameters and sharing

To characterize the intact E/c relationship, the Hill model was chosen (Gesztelyi et al, 2012):

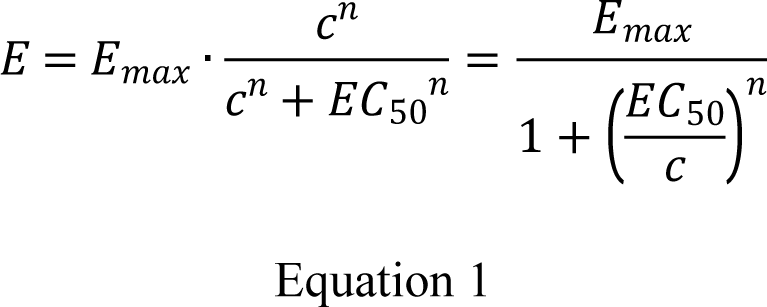

To fit the intact E/c curves, the Hill model was used in a form where all concentrations (c, EC_50_) were expressed as common logarithms (logc, logEC_50_):

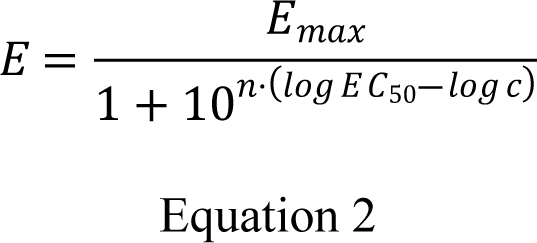

where: c: the concentration of the agonist, administered during the construction of the E/c curve; E: the effect of c (considered to be intact because of the lack of any biasing factor); E_max_: the maximal effect; EC_50_: the agonist concentration producing half-maximal effect; n: the Hill coefficient (slope factor). It should be emphasized that, during the curve fitting, logc (and not c) was the independent variable and logEC_50_ (and not EC_50_) served as a parameter (see: Appendix).

To obtain the biasing concentration, the model of RRM (see below the lower two expressions) was applied. This model was derived from the fusion of the equation 1 (see above) and the basic equation of RRM (see below the upper equation) (Gesztelyi et al, 2004; Grenczer et al, 2010a):

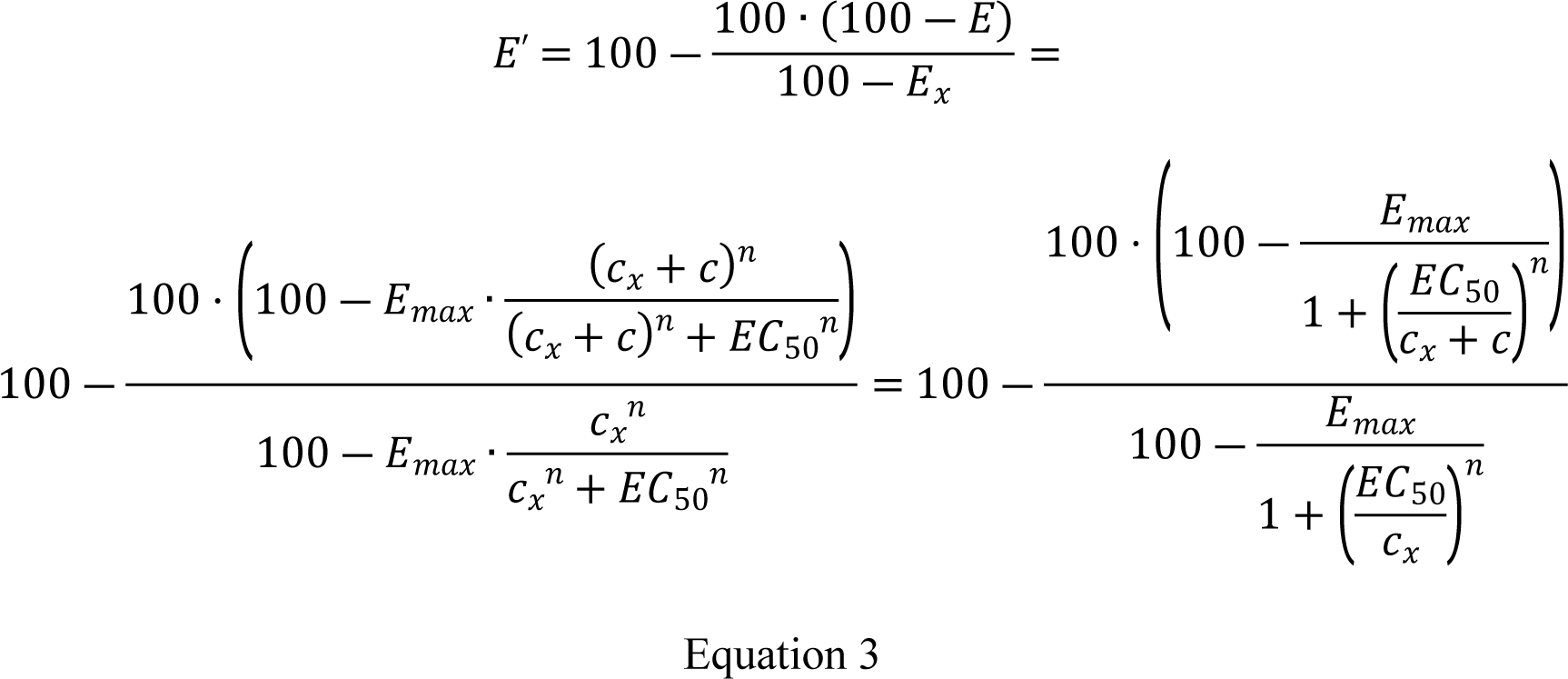

where: c: the concentration of the agonist, administered during the construction of the E/c curve; c_x_: the biasing concentration; E’: the biased effect that was calculated as the effect of c, regardless of the presence of c_x_; E: the intact counterpart of E’, i.e. the effect that properly reflects the co-action of c and c_x_; E_x_: the effect of c_x_ (that is also intact, similarly to E); E_max_, EC_50_, n: parameters identical to those with the same name in the equation 1 (indicating that the Hill model defines the relationship between agonist concentration and effect for E’, E and E_x_).

To fit both the intact and biased E/c curves, two equivalent forms of the equation 3 were used: the equation 4 that contained the common logarithm of the biasing concentration (logc_x_), and the equation 5 that included the biasing concentration itself (c_x_):

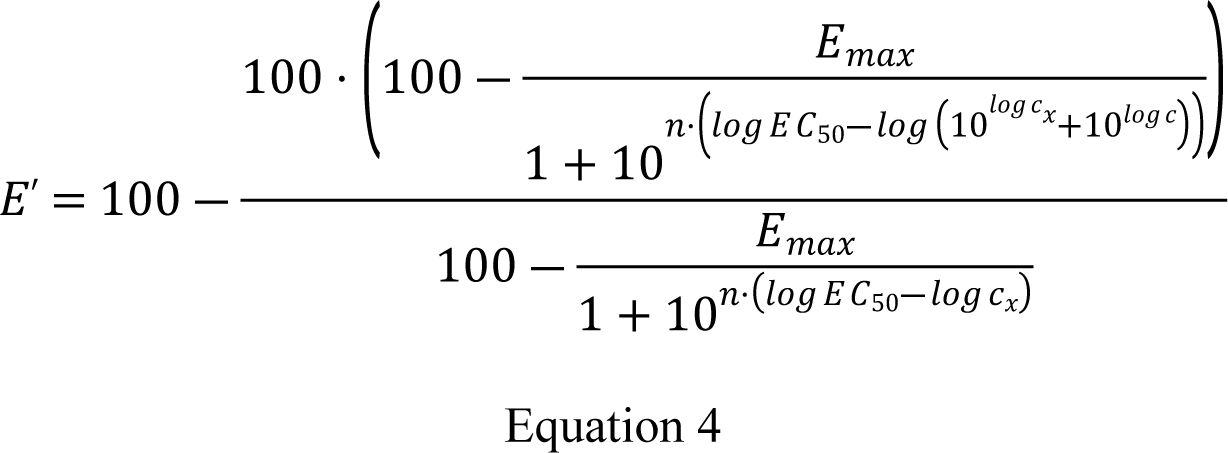

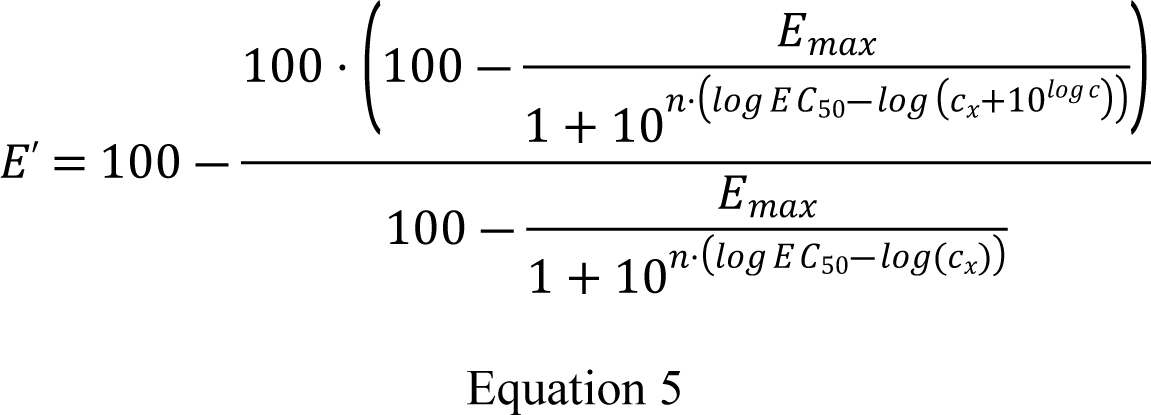

Thus, in the equation 4 and 5, logc_x_ and c_x_ served as a parameter, respectively, in addition to E_max_, logEC_50_ and n, while logc was the independent variable (see: Appendix).

Both equations 4 and 5 were fitted three ways: individually, globally with one model (i.e. only the equation 4 or 5), and globally with two models (sc. the equation 2 together with the equation 4 or 5) (see: Appendix). During individual regression, the equation 4 or 5 was fitted to one E/c curve at once (either intact or biased), in a manner that the Hill parameters (E_max_, logEC_50_ and n) were constrained to constant values that were provided by fitting the Hill model (equation 2) to the corresponding (or the same) intact E/c curve. So, during individual regression, in the equation 4 and 5 containing four parameters, there was only one variable parameter (logc_x_ and c_x_, respectively, thus parameter sharing was out of the question).

In turn, the globally fitted equation 4 or 5 included four variable parameters (E_max_, logEC_50_, n and logc_x_ or c_x_). When using one model, the related intact and biased E/c curves were simultaneously fitted to the equation 4 or 5, while the Hill parameters were shared between (or among) the E/c curves. When applying two models (the equation 2 together with the equation 4 or 5), the related intact and biased E/c curves were also simultaneously fitted with shared Hill parameters but in a way that the equation 2 was fitted exclusively to the intact E/c curve, while the equation 4 or 5 was fitted only to the related biased one(s).

When more than one biased E/c curves belonged to one intact E/c curve, there were two further options to fit globally (irrespective of the number of models): curve fitting at once to all related E/c curves (“all-at-once” manner), and curve fitting simultaneously to only two related E/c curves, one of which was the intact E/c curve (“pairwise” technique). Both options were carried out.

During the individual and one-model global ways of regression, logc_x_ and c_x_ values were obtained for the intact E/c curves as well, as an additional control (their expected value was zero). The two-model global fitting provided no estimates for the intact E/c curves because they were fitted only to the Hill model (equation 2).

### 2.3. Regression manners addressing data scatter

The above-mentioned ways of regression were combined with ordinary or robust fitting, addressing the distribution of the scatter of data points around the best-fit curve. Upon robust fitting, only accuracy (but not precision) could be compared, because robust regression prevented obtaining 95% CIs and 95% confidence and prediction bands (Curve Fitting Guide, 2023).

### 2.4. Data processing and presentation

Accuracy of the regression was characterized by the distance of the estimate from the corresponding known biasing concentration.

Precision of the regression was characterized by the width of the 95% CI of the best-fit value addressing the biasing concentration (logc_x_ or c_x_). In addition, precision of the curve fitting and precision of the E/c curve data were characterized by the distance of the 95% confidence and prediction bands, respectively, from the corresponding best-fit curve.

Further information supplied by a 95% CI was the position of the best-fit value within it. If the best-fit value was well centered (i.e. the 95% CI was symmetrical or close to it), the parameterizations of the model could be regarded appropriate (Curve Fitting Guide, 2023).

Curve plotting and fitting was performed with GraphPad Prism 9.5.1 for Windows (GraphPad Software Inc., La Jolla, CA, USA). For every setting not addressed above, the default option was used with two exceptions (see the next two paragraphs). In addition, some operations were made with Microsoft Excel for Microsoft 365 (Microsoft Co., Redmond, WA, USA).

When setting which way the software should check how well the experimental data define the model, both options, sc. “identifying ambiguous fits” and “identifying unstable parameters”, were performed (Curve Fitting Guide, 2023). The reason for this was that more estimates of the biasing concentration could be obtained with “identifying ambiguous fits”, an option dealing with the extent of parameter intertwining. So, when RRM was implemented with the default of this setting (“identifying unstable parameters”), some estimates, labelled as “ambiguous” by the former option, were missing, while the rest had practically the same values and 95% CIs.

For almost all cases, the default option for computing 95% CIs was “asymmetrical” (Curve Fitting Guide, 2023). Where not (during defining the models to fit), the “asymmetrical” option was chosen.

## 3. Results

### 3.1. Evaluation of intact – biased E/c curve pairs using models with different algebraic forms of the biasing concentration

In the case of individual fitting and two-model global regression, the models containing logc_x_ and c_x_ provided equaling estimates for the biased E/c curves that was not affected by the ordinary or robust way of fitting. The estimates for the intact E/c curves were not the same, nevertheless they showed small or close-to-zero values, as expected.

Upon one-model global regression, however, the estimates yielded by the models with logc_x_ and c_x_ differed, although to a moderate extent (regardless the use of ordinary or robust fitting). The estimates for the intact E/c curves were better (i.e. closer to zero), when the model included logc_x_ (cf. Table 1 and 2).

**Table 1.**
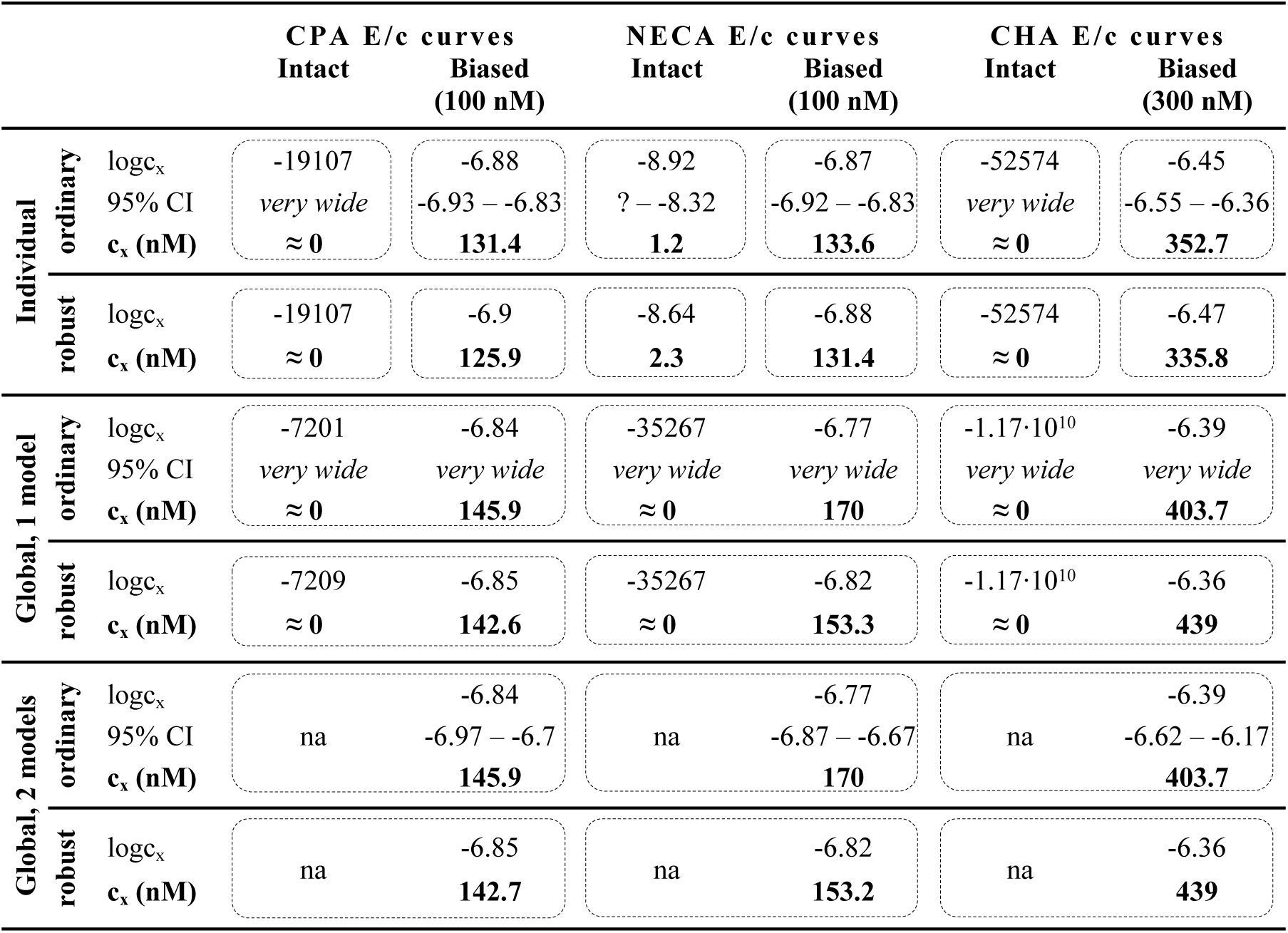
The logc_x_ best-fit values, with 95% confidence intervals (95% CI) and antilog values as estimates (c_x_, converted to nmol/L), yielded by the receptorial responsiveness method (RRM) performed on data of six groups (see column headers) of a previous *ex vivo* study (Szabo et al, 2019a), using three fitting ways combined with two another fitting options (see row headers). na: not applicable; nM: nmol/L; CPA: *N^6^*-cyclopentyladenosine; NECA: 5′-(N-ethylcarboxamido)adenosine; CHA: *N^6^*-cyclohexyladenosine

**Table 2.**
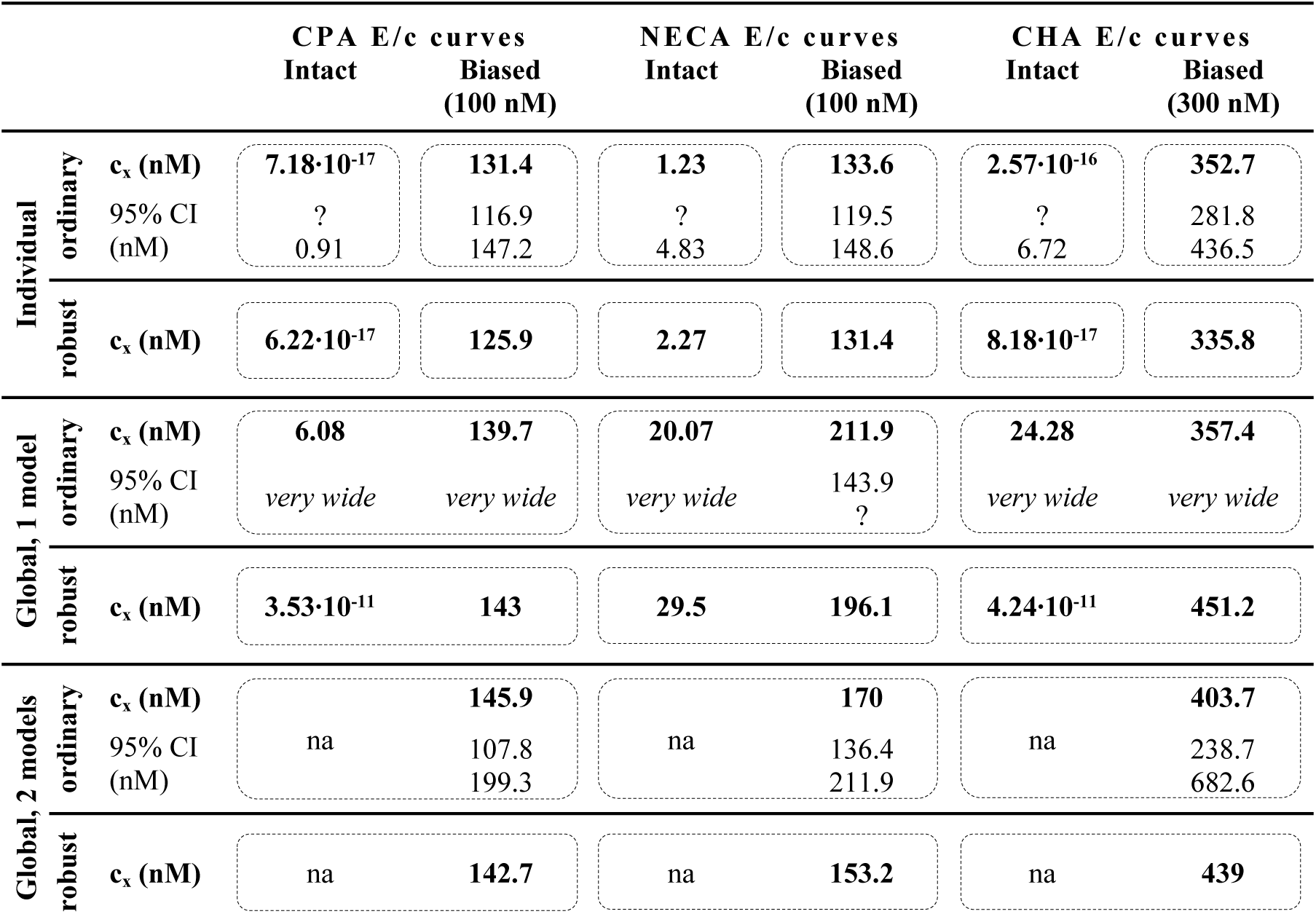
The c_x_ best-fit values (being also estimates) converted to nmol/L, with 95% confidence intervals (95% CI), provided by the receptorial responsiveness method (RRM) performed on data of six groups (see column headers) of a previous *ex vivo* study (Szabo et al, 2019a), using three fitting ways combined with two another fitting options (see row headers). na: not applicable; nM: nmol/L; CPA: *N^6^*-cyclopentyladenosine; NECA: 5′-(N-ethylcarboxamido)adenosine; CHA: *N^6^*-cyclohexyladenosine

According to the 95% CIs as well as 95% confidence and prediction bands (being accessible only upon ordinary fitting), there was no considerable difference in the precision between the models containing logc_x_ and c_x_ (cf. Table 1 and 2, furthermore the adjacent left and right panels of Fig. 1-3). As a surprising exception, when performing one-model global fitting to NECA E/c curves, confidence and prediction bands could only be plotted (i.e. only then they were not undisplayably poor) when the model contained c_x_ (rather than logc_x_) (Fig. 2).

Nevertheless, the use of c_x_ (instead of logc_x_) in the model of RRM did not generally improve either accuracy or precision of the estimation.

### 3.2. Evaluation of intact – biased E/c curve pairs with different number of models and variable parameters

Accuracy was the best in the case of individual fitting, which was followed by the other options with only moderate differences. When using the model of RRM with logc_x_, accuracy was not affected at all by the fact whether one- or two-model global fitting was chosen. Accuracy was slightly influenced by whether ordinary or robust regression was carried out, and it was moderately influenced by whether logc_x_ or c_x_ was in the fitted model (this latter observation was especially true in the case of the intact E/c curves fitted globally to one model). All estimates for the intact E/c curves showed the expected small value, especially when using logc_x_ to fit (Table 1 and 2).

According to the 95% CIs, precision was the best upon individual fitting, but only when the biased E/c curves were assessed. Precision was acceptable when using two-model global fitting (always), whereas it was unquantifiably poor in the case of one-model global regression (always) and individual fitting (when assessing intact E/c curves) (Table 1 and 2). For the one-model global regression, the cause of the ambiguous fit (or of the unstable parameters) was that logc_x_ and c_x_ were highly intertwined with the other parameters (i.e. they showed strong correlation with the other parameters). The use of c_x_ (instead of logc_x_) substantially increased the correlation even between other parameters.

The results obtained about the 95% confidence and prediction bands were supportive of those about the 95% CIs, with one exception. Namely, the 95% confidence bands for the intact E/c curves, yielded by individual fitting, seem to be surprisingly narrow, when considering the very wide 95% CIs for the corresponding best-fit values (Table 1 and 2, Fig. 1). This phenomenon was especially conspicuous when using the model with logc_x_ (Table 1, Fig. 1, left panels). This may be due to that the concerned best-fit values (both logc_x_ and c_x_) are very small (as expected in the case of intact E/c curves), so even a relatively wide range around these logc_x_ and c_x_ values can appear to be narrow. Therefore, these narrow confidence bands might be misleading, while the much wider 95% prediction bands, holding information about the uncertainty of the E/c curve data in addition to that of the curve fitting, can be considered more illustrative (Fig. 1).

Overall, the two-model global regression could be regarded the most precise, but the individual regression was not far behind, especially if considering that the estimates for the intact E/c curves served only as an additional control (Table 1 and 2, Fig. 1-3).

Regarding the convenience of use, the global way of regression was undoubtedly superior to the individual one. Specifically, the one-model global fitting was the more convenient (because it did not require creating a multiline model in the software; see: Appendix). In terms of convenience, it had no relevance whether logc_x_ or c_x_ was fitted as well as ordinary or robust regression was performed.

In addition, every best-fit value, for which 95% CI could be computed, was well centered in it, indicating the proper parametrizations of the fitted models.

### 3.3. Evaluation of E/c curve families (using only fully logarithmic models)

It is worth noting in advance that, upon global regression, the assessment of a E/c curve family offers an extra choice for the implementation of RRM: in addition to all-at-once fitting, it is possible to fit pairwise (that increases the impact of the intact E/c curve on the results). The results presented herein were obtained both ways. For simplicity, to assess E/c curve families, the model of RRM with logc_x_ was only applied.

In terms of accuracy, all manners of regression provided similarly good estimates for the greater two of the three biasing concentrations. However, the estimates for the smallest one were acceptable only when individual regression or global regression with pairwise fitting was performed. The estimates for the intact E/c curves (where relevant) were close to zero (Table 3, 4).

**Table 3.** The logc_x_ best-fit values, with 95% confidence intervals (95% CI) and antilog values as estimates (c_x_, converted to nmol/L), yielded by the receptorial responsiveness method (RRM) performed on data of four groups (see column headers) of a previous *ex vivo* study (Gesztelyi et al, 2004), using three fitting ways combined with two another fitting manners (see row headers). In the case of global regression, curve fitting was carried out by pairing each Biased group with the Intact one (“pairwise technique”). na: not applicable; nM: nmol/L; CPA: *N^6^*-cyclopentyladenosine; NECA: 5′-(N-ethylcarboxamido)adenosine; CHA: *N^6^*-cyclohexyladenosine

| Fit to curve pairs<br>(+indiv) |  |  | CPA E/c curves |  |  |  |  |  |
| --- | --- | --- | --- | --- | --- | --- | --- | --- |
|  |  |  | Intact | Biased<br>3 nM | Intact | Biased<br>30 nM | Intact | Biased<br>100 nM |
| Individual | ordinary | logc<sub>x</sub> | -14.66 | -8.1 | - | -7.58 | - | -7.02 |
|  |  | 95% CI | *very wide* | -8.2 – -8.01 | - | -7.62 – -7.55 | - | -7.09 – -6.95 |
|  |  | c<sub>x</sub> (nM) | **2.2·10<sup>-6</sup>** | **8.02** | - | **26.06** | - | **95.5** |
|  | robust | logc<sub>x</sub> | -20375 | -8.06 | - | -7.59 | - | -6.99 |
|  |  | c<sub>x</sub> (nM) | **≈ 0** | **8.66** | - | **25.46** | - | **101.4** |
| Global, 1 model | ordinary | logc<sub>x</sub> | -7237 | -8.08 | -7.04·10<sup>8</sup> | -7.54 | -1.52·10<sup>7</sup> | -6.8 |
|  |  | 95% CI | *very wide* | *very wide* | *very wide* | *very wide* | *very wide* | *very wide* |
|  |  | c<sub>x</sub> (nM) | **≈ 0** | **8.37** | **≈ 0** | **28.95** | **≈ 0** | **159.2** |
|  | robust | logc<sub>x</sub> | -7237 | -8.07 | -7.04·10<sup>8</sup> | -7.53 | -1.52·10<sup>7</sup> | -6.8 |
|  |  | c<sub>x</sub> (nM) | **≈ 0** | **8.46** | **≈ 0** | **29.83** | **≈ 0** | **158.3** |
| Global, 2 models | ordinary | logc<sub>x</sub> | na | -8.08 | na | -7.54 | na | -6.8 |
|  |  | 95% CI |  | -8.25 – -7.94 |  | -7.62 – -7.46 |  | -6.93 – -6.65 |
|  |  | c<sub>x</sub> (nM) |  | **8.37** |  | **28.95** |  | **159.2** |
|  | robust | logc<sub>x</sub> | na | -8.07 | na | -7.53 | na | -6.8 |
|  |  | c<sub>x</sub> (nM) |  | **8.46** |  | **29.83** |  | **158.3** |

**Table 4.**
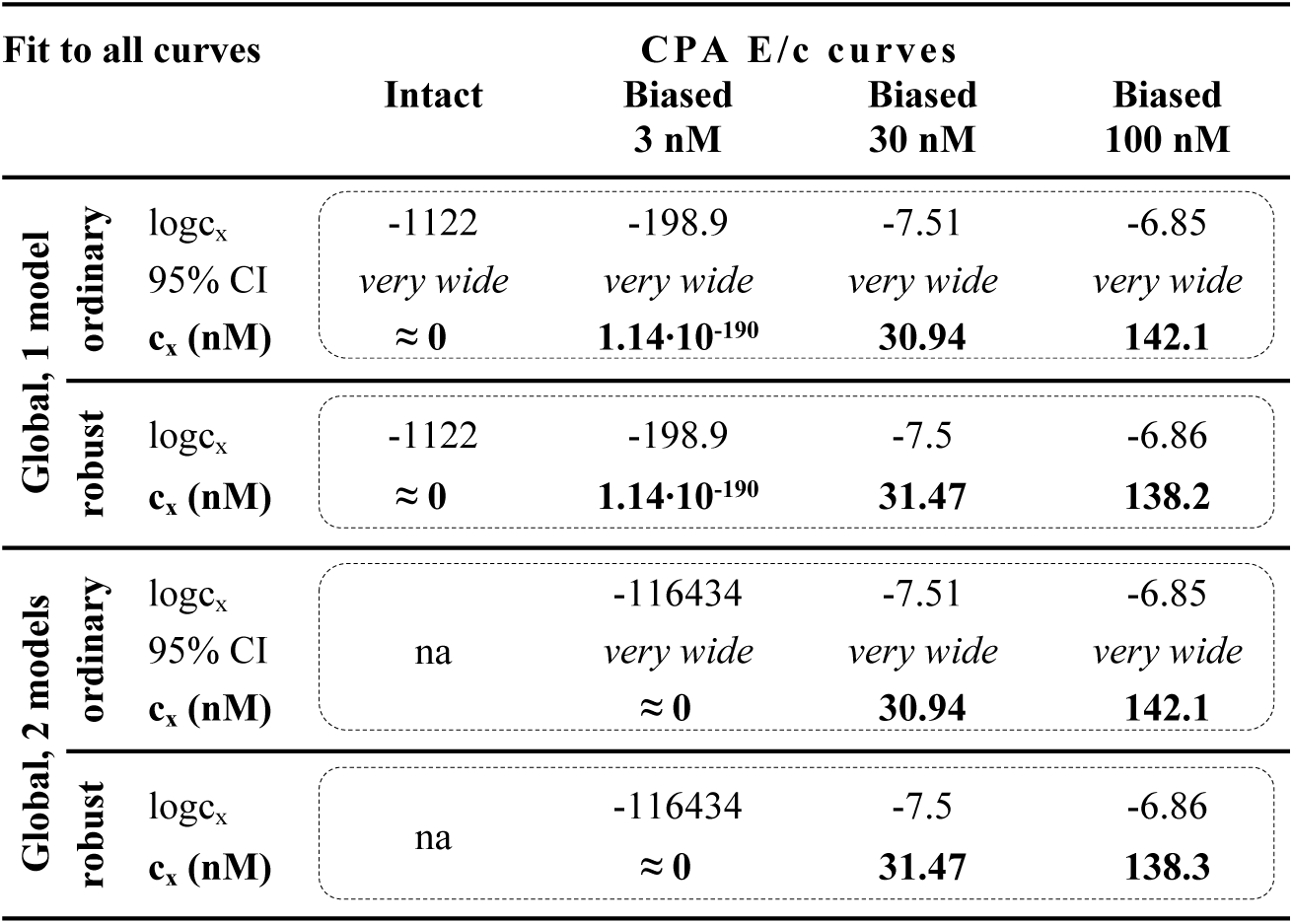
The logc_x_ best-fit values, with 95% confidence intervals (95% CI) and antilog values as estimates (c_x_, converted to nmol/L), obtained with the receptorial responsiveness method (RRM) performed on data of four groups (see column headers) of a previous *ex vivo* study (Gesztelyi et al, 2004), using two manners of global regression combined with two another fitting options (see row headers). Curve fitting was carried out simultaneously to all groups (“all-at-once technique”). na: not applicable; nM: nmol/L; CPA: *N^6^*-cyclopentyladenosine; NECA: 5′-(N-ethylcarboxamido)adenosine; CHA: *N^6^*-cyclohexyladenosine

In the case of global regression carried out with a pairwise technique, precision proved to be similar in all respects to that described in the previous subsection (for the model with logc_x_) (Table 3, 4, Fig. 4). Upon global regression performed in all-at-once manner, however, precision was unquantifiably poor, irrespective of the number of models. Thus, the two-model global regression, implemented with pairwise fitting, was considered the most precise, followed closely by the individual regression.

**Fig. 4.**
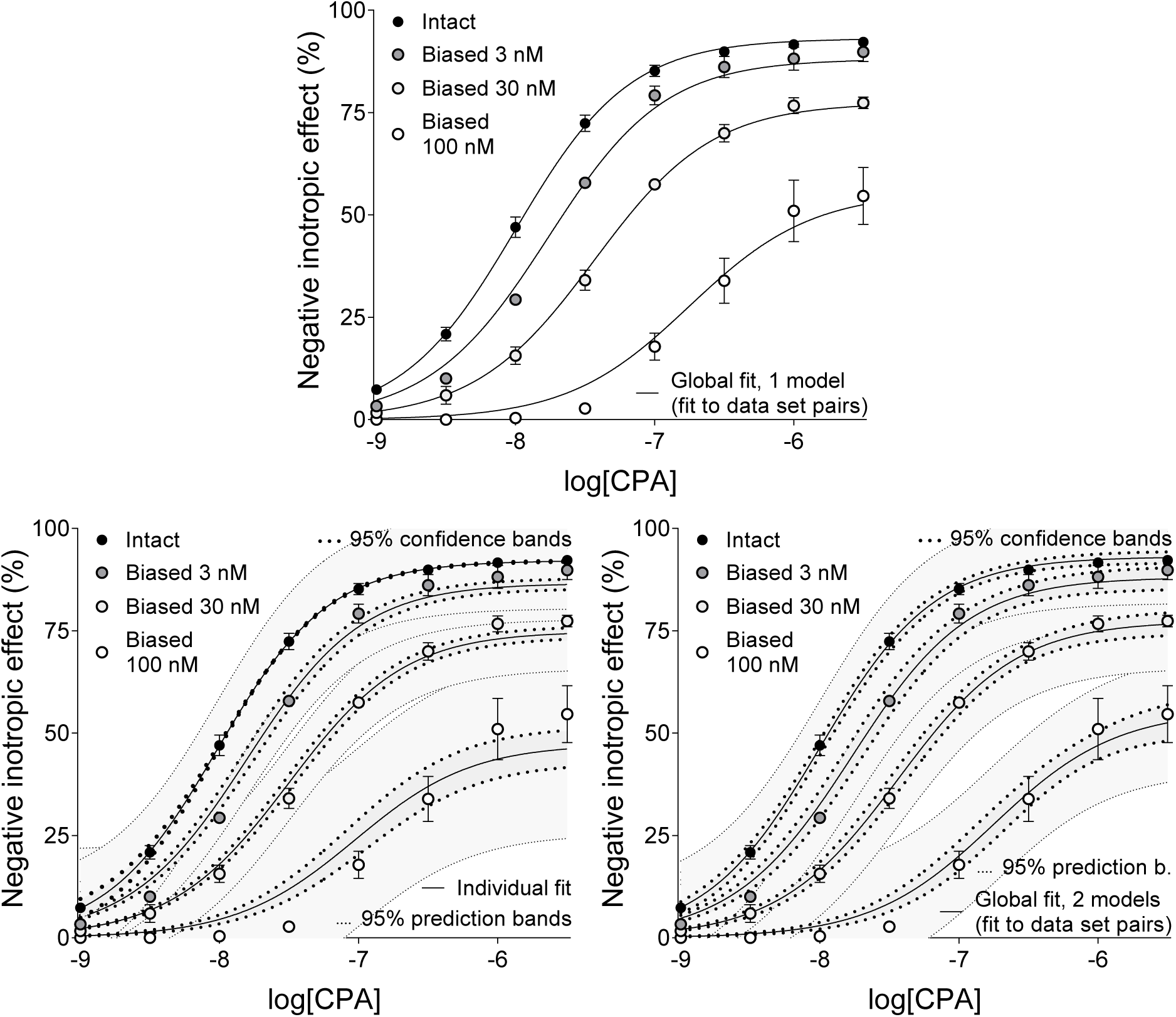
The implementation of RRM with one-model global regression (panel above), individual regression (left panel below) and two-model global regression (right panel below) combined with ordinary least-squares fitting, using exclusively the common logarithm of the concentration to be determined (logc_x_) as a parameter. The procedure was performed on concentration-effect (E/c) curve families consisting of “Intact” and “Biased” E/c curves, obtained from an earlier *ex vivo* study (Gesztelyi et al, 2004). The *x*-axis shows the common logarithm of the molar concentration of CPA, a synthetic A_1_ adenosine receptor full agonist (administered for constructing the E/c curve). The *y*-axis indicates the effect (the percentage decrease in the initial contractile force of isolated, paced guinea pig left atria). The “Intact” E/c curves (black symbols) were generated conventionally, while the “Biased” E/c curves (symbols filled with a color from dark gray to white) were constructed in the presence of a single surplus CPA concentration (which increased as the color of the symbol lightened). The symbols indicate the responses to CPA averaged within the groups (± SEM). The continuous lines show the best-fit curves of the fitted equation 2 or 4 (chosen according to the way of fitting and to the type of the E/c curve). The thick dotted lines indicate the 95% confidence bands, while the thin dotted lines denote the 95% prediction bands. RRM: receptorial responsiveness method; CPA: N^6^-cyclopentyladenosine

Regarding manageability, the one-model global regression performed with all-at-once fitting was the most convenient to use, it was followed by the one-model global regression with pairwise fitting, then the two-model global regression with all-at-once fitting, and then the two-model global regression with pairwise fitting. Eventually, the most complicated way to perform RRM was the individual regression in this case as well.

Every best-fit value was well centered in its 95% CI (if any), thus the parametrizations of the models were found to be appropriate here as well.

The main results of the study have been summarized in Table 5.

**Table 5.** Comparison of the results obtained from different implementations of the receptorial responsiveness method (RRM). These conclusions have been drawn for the model containing logc_x_ (proved to be better than that including c_x_), but roughly the same is true for the model with c_x_ as well.

|  | **Individual**<br>**(local)** | **Global** |  |  |  |
| --- | --- | --- | --- | --- | --- |
|  |  | **fit to all curves** |  | **fit to curve pairs** |  |
|  |  | **one model** | **two models** | **one model** | **two models** |
| **Accuracy** | very good | poor/good | poor/good | good | good |
| **Precision** | good | poor | poor | poor | very good |
| **Manageability** | arduous | very easy | easy | easy | complicated |

## 4. Discussion

In the present investigation, different implementations of RRM were tested by estimating known concentrations of stable adenosine receptor agonists in the microenvironment of the A_1_ adenosine receptors located in the guinea pig atrial myocardium.

RRM is a procedure based on simple nonlinear regression with a unique model that contains two variables (the concentration of an agonist, usually as a logarithm, and the effect evoked by this concentration in a biological system) plus at least one variable parameter. In the broadest sense, the role of this obligate variable parameter is to quantify something, which has caused a decrease in the virtual responsiveness of the given biological system. This “something” is usually a single, constant concentration of an agonist (“biasing agonist”), quantification of which is the typical goal of RRM. This quantification is made with the concentration of the same or another agonist that can exert the same decrease in the responsiveness as the original evoking factor (Gesztelyi et al, 2004; Grenczer et al, 2010a, 2010b).

Beyond the simple concentration estimation (such as the determination of the interstitial adenosine accumulation under nucleoside transport blockade: Karsai et al, 2006, 2007), RRM meets other types of challenges. RRM is suitable to correct E/c curves of adenosine receptor agonists for the bias caused by changes in the endogenous adenosine levels. Accordingly, RRM helped to uncover a slight positive inotropic effect, resulted partly from phosphodiesterase type 2 inhibition and partly from an unidentified mechanism, of agents with strong adenosine deaminase inhibitory effect. This latter activity led to adenosine accumulation with a consequent A_1_ adenosine receptor-mediated negative inotropy, which interfered with the positive inotropic action (Gesztelyi et al, 2003; Kemeny-Beke et al, 2007; Pak et al, 2015). RRM was also used to incriminate a hitherto unknown feature of an agent with a well-known irreversible A_1_ adenosine receptor antagonist property, namely the ability to decrease the interstitial adenosine concentration (*via* an unknown mechanism), in a series of *in silico* (Zsuga et al, 2017; Szabo et al, 2019b) and *ex vivo* investigations (Erdei et al, 2018; Viczjan et al, 2021). Furthermore, RRM contributed to the assessment of the A_1_ adenosine receptor reserve for the direct negative inotropic effect of adenosine, an agonist difficult to quantify *ex vivo* (and *in vivo*) due to its short half-life (Kiss et al, 2013; Pak et al, 2014; Zsuga et al, 2017). In addition, RRM was used to tease out two possible actions of a widely used phytocannabinoid, i.e. nucleoside transport blockade and A_1_ adenosine receptor agonism (Viczjan et al, 2022).

The model of RRM describes the link between the biased effect (measured in a system with decreased responsiveness) and the corresponding intact one (determined in the same system in its intact state), with the help of a biasing concentration (c_x_) that is attributed to an agonist that is used for the determination (see the equation 3). So, when using this model (see equations 4 and 5) for data resulted from the co-action of two different agonists, inevitably some theoretical simplification occurs.

As input for RRM, (at least) two (types of) E/c curves are needed: an “intact” E/c curve that displays the effect of the agonist used for the E/c curve in the system possessing its intact responsiveness, and a “biased” one that shows the effect of this agonist in the system with decreased response capacity. The model of RRM should be fitted to the biased E/c curve with the utilization of the information implied in the intact E/c curve (Gesztelyi et al, 2004; Grenczer et al, 2010a; Szabo et al, 2019a). The way this information is transmitted is what distinguishes individual (local) regression, one-model global regression and two-model global regression as options for implementing RRM.

A noteworthy feature of regression is that fitting of different but algebraically equivalent expressions can lead to (somewhat) different results. This is the reason for the recommendation that a parameter following log-normal (Galton) distribution (e.g. concentration) should be used as a common logarithm in models intended for curve fitting, in order to receive a symmetrical CI for it (Motulsky and Christopoulos, 2004). In the model of RRM, the biasing concentration can also be expressed as either a logarithm (logc_x_) or a numerus (c_x_), options that may affect the results of the estimation. As the software used for curve fitting in the present study can handle asymmetrical CIs as well, it seemed to be reasonable to revisit, which form of the biasing concentration is better for fitting.

The aforementioned regression manners can be further combined with two additional fitting options, ordinary and robust regression, to consider the distribution of the scatter of data points around the best-fit curve. This distribution can be closer to the Gaussian distribution or to the Lorentzian one (the two extremes of the t distribution regarding the number of degrees of freedom), requiring ordinary or robust regression, respectively. It should be noted that the use of robust regression limits the usefulness of the estimation as it prevents the determination of precision (Curve Fitting Guide, 2023).

It should also be mentioned that RRM can handle if more than one biased E/c curve belong to one intact E/c curve (Gesztelyi et al, 2004). However, when choosing global regression, this circumstance raises two new possibilities to perform RRM: fitting all related E/c curves at once or fitting them pairwise, i.e. each biased E/c curve separately with the intact E/c curve.

Thus, regression, the essence of RRM, may be carried out several ways. The goal of this study was to find the best from these possibilities. On one hand, the individual regression (the first and, up to now, the most used manner to implement RRM), the one-model global regression (proved to be more convenient but less accurate and precise than the individual one: Szabo et al, 2019a), and the two-model global regression (an untried possibility yet) were compared. Furthermore, these choices were combined with other options, i.e. fitting the parameter estimating the biasing concentration as a logarithm (logc_x_) or a numerus (c_x_), performing ordinary or robust regression, furthermore applying all-at-once or pairwise fitting techniques (where appropriate).

In the present study, accuracy of RRM have generally been found to be acceptable for all kinds of regression used here (Table 1-4). Nevertheless, there were two major exceptions: the intact E/c curves determined with one-model global regression using a model containing c_x_ (Table 2), and the smallest biasing concentration measured with global fitting in an all-at-once way (Table 4). It can be concluded that a small biasing concentration, the effect of which (E_x_ in the equation 3) is small relative to the maximal effect of the agonist used for the E/c curves (E_max_ in the equation 3), poses a challenge for RRM. This is consistent with our earlier finding that, using RRM, too small and too great concentrations may be estimated with substantial inaccuracy (Grenczer et al, 2010a, 2010b).

It can be concluded that most of the differences among the various implementations of RRM have been found in terms of precision and convenience of use. Accordingly, choosing between the ordinary or robust way of regression had little influence on the present results, as the robust regression prevented calculating confidence intervals for the best-fit values as well as confidence and prediction bands for the best-fit curves. Besides, the robust regression did not affect the convenience (manageability) of curve fitting at all (Table 1-4).

In addition, it has been observed that rewriting the RRM’s model to replace logc_x_ with c_x_ did not improve either accuracy or precision of the estimation. Moreover, in some cases, the use of c_x_ worsened precision by increasing the correlation between some parameters. This finding has disproved our assumption that the poor performance of global regression (using one model) in a previous investigation (Szabo et al, 2019a) might be due to the logarithmic form of the parameter estimating the biasing concentration (logc_x_ instead of c_x_). Thus, there is no reason to replace the fully logarithmic model of RRM with one that contains c_x_ as a parameter.

Furthermore, for E/c curve families, accuracy (when a small concentration was to be determined) and precision (in every case) of the global regression could be substantially improved by fitting the appropriate E/c curves pairwise (cf. Table 3 and 4). Thus, despite the proper estimates for most cases and the ease of use, we do not recommend the global fitting simultaneously to more than two E/c curves, when performing RRM. In line with this, we discuss hereafter only those results of global regression that were obtained with pairwise fitting technique (unless otherwise indicated).

Overall, in terms of accuracy, the individual fitting proved to be the best, followed closely by the global regression (when fitting the model with logc_x_, irrespective of the number of models). In turn, regarding precision (i.e. the reliability of the estimates), the two-model global fitting was the best, followed closely by the individual fitting and, from afar, by the one-model global fitting (Table 1-4). Regarding manageability, the one-model global regression was the easiest to use, followed by the moderately complicated two-model global fitting, and then by the most complicated individual regression (Table 5).

The poor precision of the one-model global regression has been indicated by the fact that (upon ordinary fitting) neither confidence intervals nor confidence and prediction bands could be obtained for the results (interestingly, except for the NECA E/c curves when fitting c_x_ as a parameter) (Fig. 2). The source of this uncertainty was that the parameter estimating the biasing concentration (logc_x_ or c_x_) was greatly dependent on the other parameters when fitting the intact E/c curves. The other occasion, when similarly high level of parameter correlation occurred, was the attempt to determine the smallest biasing concentration by fitting simultaneously more than two E/c curves (affecting both types of global regression).

In favor of the oldest way to perform RRM, it should be noted that the individual regression imposes the fewest requirements for the curve fitting software used. If it is not possible to fix the Hill parameters at constant values, then these values can simply be written into the model of RRM, and the model individualized this way can be fitted. Fortunately, there are nowadays much software with great capabilities to choose from.

Consequently, neither the RRM’s model with c_x_ (regarding both accuracy and precision) nor the two-model global fitting (in terms of accuracy) exhibited a smashing advantage over the well-established individual regression using a fully logarithmic model. This indicates that the foibles of RRM in any case do not stem from the fit of logc_x_ that would hinder the correct determination of the zero value of c_x_ for the intact E/c curves. So, the problems observed in some cases may be the results of the relative complexity of the model. The greater the complexity of a regression model, the greater the degree of correlation that can occur between the parameters of the model (Motulsky and Christopoulos, 2004; Curve Fitting Guide, 2023). In cases when the agonist of the E/c curves and the biasing agonist are different, the causal role of the theoretical simplification occurring during the use of RRM may also be significant in terms of accuracy as well as precision (Grenczer et al, 2010a, 2010b).

Taking all together, for the implementation of RRM, we recommend the two-model global regression, performed with a pairwise technique (i.e. fitting only two E/c curves at once, from which one is always the intact E/c curve), when appropriate. Nevertheless, the individual regression is a good alternative to the two-model global regression. For data with too much scatter, choosing robust fitting (instead of the first-choice ordinary least-squares regression) may be considered.

## Data availability

Regarding data used for the present work, any inquiries can be directed to the corresponding author.

## Appendix

In this section, the variables and parameters have been marked as they were used in the curve fitting software. For example, the independent and dependent variables are X and Y here, while elsewhere they are logc and E (or E’), respectively.

As Hill model, the software’s built-in equation “sigmoidal dose-response (variable slope)”, with a “bottom” parameter constrained to zero (Curve Fitting Guide, 2023), was used (equivalent with the equation 2):

**Y = Emax/(1+10^(n*(logEC50-X)))**

The models of RRM written into the software for the individual and one-model global fitting; when using the logarithm of c_x_ (equivalent with the equation 4):

**Y = 100-100*(100-Emax/(1+10^(n*(logEC50-log(10^logcx+10^X)))))/(100-Emax/(1+10^(n*(logEC50-logcx))))**

and when using the c_x_ itself (equivalent with the equation 5):

**Y = 100-100*(100-Emax/(1+10^(n*(logEC50-log(cx+10^X)))))/(100-Emax/(1+10^(n*(logEC50-log(cx)))))**

The multiline model written into the software for the two-model global fitting; when using the logarithm of c_x_ (see equations 2 and 4):

**Hill = Emax/(1+10^(n*(logEC50-X)))**

**RRM = 100-100*(100-Emax/(1+10^(n*(logEC50-log(10^logcx+10^X)))))/(100-Emax/(1+10^(n*(logEC50-logcx))))**

**<A> Y = Hill**

**<∼A> Y = RRM**

and when applying the c_x_ itself (see equations 2 and 5):

**Hill = Emax/(1+10^(n*(logEC50-X)))**

**RRM = 100-100*(100-Emax/(1+10^(n*(logEC50-log(cx+10^X)))))/(100-**

**Emax/(1+10^(n*(logEC50-log(cx)))))**

**<A> Y = Hill**

**<∼A> Y = RRM**

(For more detail, see “fitting different models to different data sets” in: Curve Fitting Guide, 2023.)

## References

1. Anjos RSD, Nóbrega RS, Ferreira HDS, Lacerda AP, Sousa-Neves N. Exploring local and global regression models to estimate the spatial variability of Zika and Chikungunya cases in Recife, Brazil. Rev Soc Bras Med Trop. 2020; 53: e20200027. doi: 10.1590/0037-8682-0027-2020

2. Curve Fitting Guide. https://www.graphpad.com/guides/prism/latest/curve-fitting/index.htm (accessed on 18 March 2023)

3. Erdei T, Szabo AM, Lampe N, Szabo K, Kiss R, Zsuga J, Papp C, Pinter A, Szentmiklosi AJ, Szilvassy Z, Juhasz B, Gesztelyi R. FSCPX, a chemical widely used as an irreversible A1 adenosine receptor antagonist, modifies the effect of NBTI, a nucleoside transport inhibitor, by reducing the interstitial adenosine level in the guinea pig atrium. Molecules. 2018; 23: E2186. doi: 10.3390/molecules23092186

4. Gesztelyi R, Zsuga J, Hajdú P, Szabó JZ, Cseppento A, Szentmiklósi AJ. Positive inotropic effect of the inhibition of cyclic GMP-stimulated 3’,5’-cyclic nucleotide phosphodiesterase (PDE2) on guinea pig left atria in eu- and hyperthyroidism. Gen Physiol Biophys. 2003; 22: 501–13.

5. Gesztelyi R, Zsuga J, Juhász B, Dér P, Vecsernyés M, Szentmiklósi AJ. Concentration estimation via curve fitting: quantification of negative inotropic agents by using a simple mathematical method in guinea pig atria. Bull Math Biol. 2004; 66: 1439–53. doi: 10.1016/j.bulm.2004.03.001

6. Gesztelyi R, Zsuga J, Kemeny-Beke A, Varga B, Juhász B, Tosaki A. The Hill equation and the origin of quantitative pharmacology. Arch Hist Exact Sci. 2012; 66: 427–438, doi:10.1007/s00407-012-0098-5

7. Grenczer M, Pinter A, Zsuga J, Kemeny-Beke A, Juhasz B, Szodoray P, Tosaki A, Gesztelyi R. The influence of affinity, efficacy, and slope factor on the estimates obtained by the receptorial responsiveness method (RRM): a computer simulation study. Can J Physiol Pharmacol. 2010a; 88: 1061–73. doi: 10.1139/y10-078

8. Grenczer M, Zsuga J, Majoros L, Pinter A, Kemeny-Beke A, Juhasz B, Tosaki A, Gesztelyi R. Effect of asymmetry of concentration-response curves on the results obtained by the receptorial responsiveness method (RRM): an in silico study. Can J Physiol Pharmacol. 2010b; 88: 1074–83. doi: 10.1139/y10-089

9. Herman P, Lee JC. The Advantage of Global Fitting of Data Involving Complex Linked Reactions. In: Fenton A. (ed) Allostery. Methods in Molecular Biology. Springer, New York. 2012. doi: 10.1007/978-1-61779-334-9_22

10. Karsai D, Gesztelyi R, Zsuga J, Jakab A, Szendrei L, Juhasz B, Bak I, Szabo G, Lekli I, Vecsernyes M, Varga E, Szentmiklosi AJ, Tosaki A. Influence of hyperthyroidism on the effect of adenosine transport blockade assessed by a novel method in guinea pig atria. Cell Biochem Biophys. 2007; 47: 45–52. doi: 10.1385/cbb:47:1:45

11. Karsai D, Zsuga J, Juhasz B, Der P, Szentmiklosi AJ, Tosaki A, Gesztelyi R. Effect of nucleoside transport blockade on the interstitial adenosine level characterized by a novel method in guinea pig atria. J Cardiovasc Pharmacol. 2006; 47: 103–9. doi: 10.1097/01.fjc.0000196239.51018.a0

12. Kemeny-Beke A, Jakab A, Zsuga J, Vecsernyes M, Karsai D, Pasztor F, Grenczer M, Szentmiklosi AJ, Berta A, Gesztelyi R. Adenosine deaminase inhibition enhances the inotropic response mediated by A1 adenosine receptor in hyperthyroid guinea pig atrium. Pharmacol Res. 2007; 56: 124–31.

13. Kiss Z, Pak K, Zsuga J, Juhasz B, Varga B, Szentmiklosi AJ, Haines DD, Tosaki A, Gesztelyi R. The guinea pig atrial A1 adenosine receptor reserve for the direct negative inotropic effect of adenosine. Gen Physiol Biophys. 2013; 32: 325–35. doi: 10.4149/gpb_2013041

14. Klaasse EC, Ijzerman AP, de Grip WJ, Beukers MW. Internalization and desensitization of adenosine receptors. Purinergic Signal. 2008; 4: 21–37. doi: 10.1007/s11302-007-9086-7

15. Knutson JR, Beechem JM, Brand L. Simultaneous analysis of multiple fluorescence decay curves: A global approach. Chem Phys Lett. 1983; 102: 501–507. doi: 10.1016/0009-2614(83)87454-5

16. Motulsky HJ, Christopoulos A. Fitting models to biological data using linear and nonlinear regression. A practical guide to curve fitting. GraphPad Software Inc., San Diego, 2004. https://cdn.graphpad.com/faq/2/file/Prism_v4_Fitting_Models_to_Biological_Data.pdf (accessed on 8 April 2023)

17. Mundell S, Kelly E. Adenosine receptor desensitization and trafficking. Biochim Biophys Acta. 2011; 1808: 1319-1328. doi:10.1016/j.bbamem.2010.06.007

18. Pak K, Papp C, Galajda Z, Szerafin T, Varga B, Juhasz B, Haines D, Szentmiklosi AJ, Tosaki A, Gesztelyi R. Approximation of A1 adenosine receptor reserve appertaining to the direct negative inotropic effect of adenosine in hyperthyroid guinea pig left atria. Gen Physiol Biophys. 2014; 33: 177–88. doi: 10.4149/gpb_2013079

19. Pak K, Zsuga J, Kepes Z, Erdei T, Varga B, Juhasz B, Szentmiklosi AJ, Gesztelyi R. The effect of adenosine deaminase inhibition on the A1 adenosinergic and M2 muscarinergic control of contractility in eu- and hyperthyroid guinea pig atria. Naunyn Schmiedebergs Arch Pharmacol. 2015; 388: 853–68. doi: 10.1007/s00210-015-1121-6

20. Ramakers BP, Pickkers P, Deussen A, Rongen GA, van den Broek P, van der Hoeven JG, Smits P, Riksen NP. Measurement of the endogenous adenosine concentration in humans in vivo: methodological considerations. Curr Drug Metab. 2008; 9: 679–85. doi: 10.2174/138920008786049249

21. Spatial Data Science: https://rspatial.org/analysis/6-local_regression.html (accessed on 9 May 2023)

22. Szabo AM, Erdei T, Viczjan G, Kiss R, Zsuga J, Papp C, Pinter A, Juhasz B, Szilvassy Z, Gesztelyi R. An advanced in silico modelling of the interaction between FSCPX, an irreversible A1 adenosine receptor antagonist, and NBTI, a nucleoside transport inhibitor, in the guinea pig atrium. Molecules. 2019b; 24: E2207. doi: 10.3390/molecules24122207

23. Szabo AM, Viczjan G, Erdei T, Simon I, Kiss R, Szentmiklosi AJ, Juhasz B, Papp C, Zsuga J, Pinter A, Szilvassy Z, Gesztelyi R. Accuracy and Precision of the Receptorial Responsiveness Method (RRM) in the Quantification of A1 Adenosine Receptor Agonists. Int J Mol Sci. 2019a; 20: 6264. doi: 10.3390/ijms20246264

24. Valcárcel M. The Analytical Problem. In: Principles of Analytical Chemistry. Springer, Berlin, Heidelberg. 2000. doi: 10.1007/978-3-642-57157-2_7

25. Viczjan G, Erdei T, Ovari I, Lampe N, Szekeres R, Bombicz M, Takacs B, Szilagyi A, Zsuga J, Szilvassy Z, Juhasz B, Gesztelyi R. A body of circumstantial evidence for the irreversible ectonucleotidase inhibitory action of FSCPX, an agent known as a selective irreversible A1 adenosine receptor antagonist so far. Int J Mol Sci. 2021; 22: 9831. doi: 10.3390/ijms22189831

26. Viczjan G, Szilagyi A, Takacs B, Ovari I, Szekeres R, Tarjanyi V, Erdei T, Teleki V, Zsuga J, Szilvassy Z, Juhasz B, Varga B, Gesztelyi R. The effect of a long-term treatment with cannabidiol-rich hemp extract oil on the adenosinergic system of the Zucker diabetic fatty (ZDF) rat atrium. Front Pharmacol. 2022; 13: 1043275. doi: 10.3389/fphar.2022.1043275

27. Willems L, Reichelt ME, Molina JG, Sun CX, Chunn JL, Ashton KJ, Schnermann J, Blackburn MR, Headrick JP. Effects of adenosine deaminase and A1 receptor deficiency in normoxic and ischaemic mouse hearts. Cardiovasc Res. 2006; 71: 79–87. doi:10.1016/j.cardiores.2006.03.006

28. Zsuga J, Erdei T, Szabó K, Lampe N, Papp C, Pinter A, Szentmiklosi AJ, Juhasz B, Szilvássy Z, Gesztelyi R. Methodical challenges and a possible resolution in the assessment of receptor reserve for adenosine, an agonist with short half-life. Molecules. 2017; 22: E839. doi: 10.3390/molecules22050839

